# End-repair causes methylation underestimation in cell-free DNA sequencing libraries

**DOI:** 10.64898/2025.12.15.694439

**Authors:** Tobin E. Groth, Andrew A. Mishin, Varsha Rao, Ruby Tibet, Christopher J. Troll

**Affiliations:** Claret Bioscience LLC, 100 Enterprise Way, Suite A102, Scotts Valley, CA 95066, USA

**Keywords:** Cell-free DNA (cfDNA), Methylation sequencing, End-repair, Single-stranded DNA (ssDNA) library preparation, Double-stranded DNA (dsDNA) library preparation, Fragmentomics, Nucleosome positioning, Tissue of origin (TOO)

## Abstract

Cell-free DNA methylation sequencing provides insight into tissue of origin and chromatin structure. In some workflows, generating libraries includes end-repair. Using matched single-stranded and double-stranded libraries prepared from the same cfDNA extracts, we show that end-repair in double-stranded DNA libraries reduces globally inferred CpG methylation leading to decreased tissue of origin accuracy. Trimming read termini partially mitigates this bias but decreases coverage and removes fragmentomic information compared to single-stranded DNA libraries, which forego end-repair.

## Manuscript

### Introduction

Cell-free DNA (cfDNA) in plasma reflects ongoing cellular turnover and provides a non-invasive window into tissue biology[1–3]. cfDNA fragmentation is non-random: nucleosomes protect DNA from nuclease digestion while progressive unwrapping exposes minor grooves to nucleases such as DNASE1L3, producing phased cuts and generating the canonical mononucleosome peak with its ∼10 bp sawtooth substructure [4,5]. cfDNA molecules are preferentially derived from nucleosome-protected regions of closed chromatin, which are less accessible to endonuclease cleavage and are typically more heavily methylated than open chromatin[4,6,7]. As a result, cfDNA is globally enriched for methylation relative to genomic DNA. Consequently, methylation sequencing of cfDNA provides rich epigenomic information that reflects tissue composition, cellular turnover, and chromatin state across physiological and disease contexts [6,8,9].

Fragmentomic analyses of cfDNA termini have revealed a high frequency of longer 5’ and 3’ overhangs compared with sonicated genomic DNA[10]. During typical double-stranded DNA (dsDNA) ligation based library preparation methods, these irregular ends are converted to blunt ends through enzymatic end-repair, a step that replaces 3’ recessed bases with newly synthesized, unmethylated nucleotides. This process lowers CpG methylation near fragment termini[10,11]. In contrast, many single-stranded DNA (ssDNA) library preparations omit end-repair and ligate adapters directly to the native DNA fragment termini, preserving original termini sequence and fragment length[12,13]. Previous studies using cfDNA have shown that library preparation method influences global methylation levels and tissue-of-origin assignment[11], but how end repair interacts with nucleosome-phased cfDNA fragmentation and affects tissue-of-origin (TOO) inference has not been systematically examined. Here we directly compare matched ssDNA and dsDNA cfDNA libraries to quantify how end-repair alters methylation patterns and downstream biological interpretation.

### Methods

Blood plasma was extracted from whole blood using the QIAamp Circulating Cell-free DNA kit (Qiagen Technologies) following the manufacturer’s protocol. The purified cfDNA was measured using a Qubit^®^ Fluorometer and an Agilent TapeStation^®^ (D5000 or D1000 High Sensitivity reagents).

All libraries were prepared from 1 ng of cfDNA input. dsDNA libraries were generated using the NEBNext® Enzymatic Methyl-seq v2 Conversion Kit following the manufacturer’s instructions and amplified for 11 cycles using NEBNext® Q5U Master Mix and in-house indexes. ssDNA libraries were prepared using the Claret Bioscience SRSLY® PicoPlus kit: cfDNA was denatured, adapters containing 5mC-protected bases were ligated, then libraries were methyl-converted using the NEBNext® Enzymatic Methyl-seq v2 Conversion Module prior to 11 cycles of amplification with the same reagents. Final molarities were estimated from fragment length distributions and DNA concentrations (Agilent TapeStation 4200 and Qubit Fluorometer). All libraries were pooled and sequenced on an Illumina NextSeq® 2000 P3 flow cell (2 × 66 bp reads).

Following sequencing, adapters were trimmed using SeqPrep2 (default parameters)[14]. Once adapters were trimmed, the reads were mapped to hg38 using bwa-meth. Duplicates in the resulting BAM file were marked using Picard’s MarkDuplicates with default parameters except for VALIDATION_STRINGENCY = LENIENT and OPTICAL_DUPLICATE_PIXEL_DISTANCE = 2500 due to the NextSeq 2000 patterned flow cell [15]. CpG site information was extracted from the BAM file using MethylDackel extract, with a minimum depth set to 3 and the minimum map quality (MAPQ) set to 20. bedtools intersect was used to subset the BED files [16]. To determine methylation of CpG sites across the read, MethylDackel’s mbias was used with the minimum MAPQ of 20.

To compare methylation relative to nucleosome position, a nucleosome map from NucPosDB/Song et al.[17] was modified to represent 1 bp midpoints instead of 100 bp regions. CpG site distances to these midpoints were calculated using bedtools closest (-D ref), and a custom script summarized average methylation as a function of distance from the nucleosome center. For dyad-specific analyses, reads were subset by strand orientation using samtools view (-L). MethylDackel extract (minimum depth 3, MAPQ 20) was used to quantify CpG methylation, which was integrated with the midpoint data using bedtools closest and summarized with custom scripts.

Tissue methylation data were obtained from Loyfer et al. [18], focusing on cfDNA-contributing tissues (vascular endothelium, lymphocytes, monocytes, granulocytes, hepatocytes, and erythrocyte progenitors). Tissue bigWig files (hg38) were downloaded from GEO, converted to bedGraph with bigWigToBedGraph, filtered to remove invalid entries, and adjusted to 1 bp resolution to match sample CpG coordinates. CpG sites were subset using two approaches: (1) filtering tissue bedGraphs by methylation fraction thresholds and (2) incorporating replicate-validated unmethylated controls from Loyfer et al. Supplementary Data 2. Tissue-specific bed files were merged, converted from hg19 to hg38 with liftOver, and intersected with sample libraries using bedtools to build CpG matrices (rows = sites, columns = tissues). Tissue-of-origin (TOO) concordance was estimated by non-negative least squares (NNLS) implemented in Python’s SciPy, using 70% of CpG sites for training and 30% for testing. The resulting predictions were evaluated with scikit-learn’s R^2^ and mean-squared-error (MSE) functions across 100 random train/test iterations.

To assess whether trimming improved tissue concordance, R2 reads were truncated by 20 or 40 bp using cutadapt (-u), and read pairs with completely trimmed R2 sequences were removed with bbmap’s reformat.sh. Trimmed R1 and R2 reads were reprocessed through alignment and methylation extraction as described above. BAM files were downsampled with samtools to the depth of the lowest-coverage sample, and CpG methylation was re-extracted using MethylDackel (minimum coverage 3, MAPQ 20). Shared CpG sites were identified with bedtools intersect, and these subsets were used for NNLS-based tissue-of-origin analysis following the same procedure outlined previously.

All figures were created in RStudio version 2024.12.0.

### Results

To examine how library preparation influences methylation signals in cfDNA, we prepared both ssDNA and dsDNA libraries from the same cfDNA extracts in replicate (Fig. 1; Supplementary Table 1). A simplified schematic overview of the two workflows is shown in Fig. 1A (see methods for full protocol). dsDNA libraries exhibit progressive losses of CpG methylation across the read, beginning as early as ∼40 bp into the forward template read and culminating in a sharp terminal decline at the 3’ end of reverse template read, whereas ssDNA libraries maintain uniform methylation across both reads (Fig. 1B). When CpGs are aligned by read orientation relative to nucleosome dyads, dsDNA libraries display hypomethylation downstream of the dyad for both forward and reverse map reads. This is consistent with methylation loss introduced during end-repair filling in 5’ overhangs. In contrast, ssDNA libraries maintain a symmetric methylation pattern around the dyad. (Fig. 1C,D). Collectively, end repair leads to a global reduction of approximately 5–7% CpG methylation across nucleosome-phased regions (Fig. 1E), superimposed on the canonical dyad-centered methylation dip shared by both library types due to restricted methyltransferase access within the nucleosome core[19,20].

**Figure 1:**
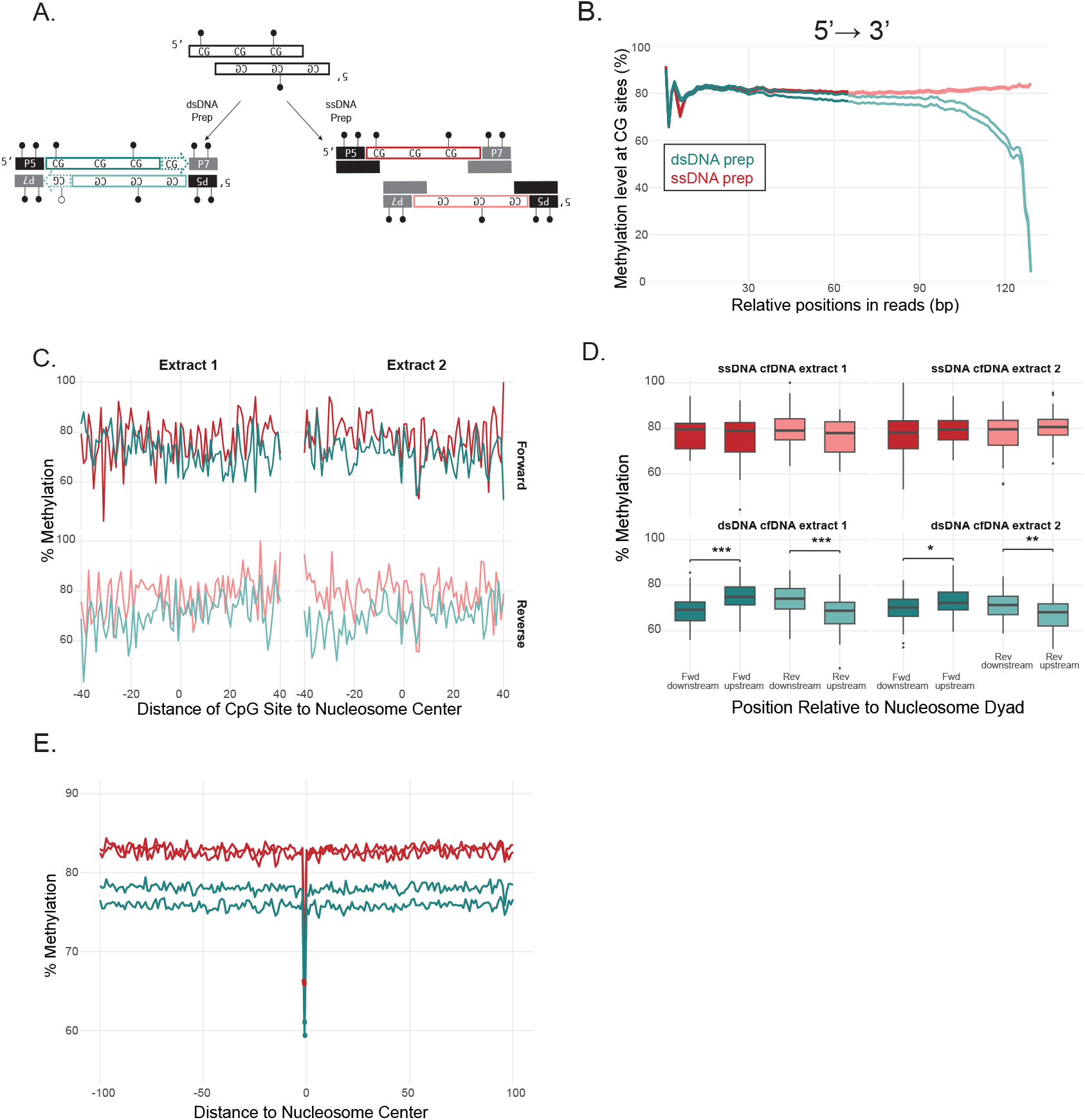
Library preparation influences cfDNA methylation profiles. A. Abbreviated schematic of ssDNA and dsDNA library preparation workflows showcasing fragments with 5’ overhangs. dsDNA libraries include an enzymatic end repair step that fills 5’ overhangs prior to methyl-protected adapter ligation and subsequent methyl-conversion. ssDNA libraries omit end repair and preserve native termini. For simplicity, individual protocol steps grouped together; methyl-conversion and index PCR not shown. B. CpG methylation along sequencing reads. Forward template (F) and reverse template (R) mapped reads (q20 and above) plotted sequentially for each library type in 5’ ⟶ 3’ direction. X-axis shows read position, y-axis shows mean CpG methylation across all covered sites. C. Strand- and position-specific CpG methylation relative to nucleosome dyads. Forward- and reverse-mapping reads are plotted separately, with distance to dyad on the x-axis and mean CpG methylation on the y-axis. D. Boxplots comparing CpG methylation upstream versus downstream of nucleosome dyads for each library type and read orientation. Each box summarizes methylation values from all CpGs in that category; Wilcoxon p-values assess differences between upstream and downstream groups. E. CpG methylation relative to nucleosome dyad centers (±100 bp). Lines show average methylation profiles centered on dyad midpoints.

We next asked whether the read-level and nucleosome-aligned methylation changes introduced by end-repair in dsDNA libraries affect TOO inference. Collapsing CpG methylation density distributions reveals that ssDNA libraries concentrate signal at fully methylated sites, consistent with the preservation of hypermethylated, nucleosome-protected regions of closed chromatin. In contrast dsDNA libraries show an inflated abundance of partially methylated sites (Fig. 2A). This redistribution results directly from the incorporation of unmethylated cytosines during end-repair, leading dsDNA libraries to underrepresent the highly methylated CpGs that are informative for tissue-of-origin deconvolution. A representative IGV view illustrates the same effect at the single-locus level, where ssDNA reads preserve methylation calls while dsDNA reads display losses at fragment 3’ termini (Fig. 2B). To quantify the impact of these end-repair–driven methylation changes on TOO analysis, we performed 100 iterations of non-negative least-squares (NNLS) analysis using a plasma-relevant atlas of immune and parenchymal cell types (granulocytes, monocytes, CD4 and CD8 T cells, liver hepatocytes, kidney epithelium, and lung endothelium; Supplementary Table 2) from Loyfer *et al*.[18]. For each iteration, 70% of CpG sites were used for training and 30% for testing to calculate R^2^ and mean-squared error (MSE) using scikit-learn. Across both cfDNA extracts, ssDNA libraries yield consistently higher R^2^ and lower MSE than dsDNA libraries for informative CpG subsets (<0.2 or >0.8 and <0.1 or >0.9 fraction methylation thresholds) (Fig. 2C–D). In contrast, the atlas-validated unmethylated control CpG set, sites that remain consistently unmethylated across tissues and replicates, shows nearly identical performance between library types, confirming that these effects are restricted to methylated, biologically informative regions. Previous work with methyl-seq on genomic DNA has shown that trimming several bases from the 3’ end of read 2 can reduce end-repair–associated methylation bias and improve signal accuracy. To test whether end-repair related methylation bias in cfDNA could be corrected in a similar approach, we progressively trimmed 20 bp and 40 bp from dsDNA reads. Trimming increased global methylation levels toward those observed in ssDNA libraries (Fig. 2E) and improved tissue-of-origin concordance and error metrics (Fig. 2F; Supplementary Fig. 1). Trimming 40 bp largely restores dsDNA methylation fidelity when ssDNA data were downsampled to similar read depth. However, this approach reduced unique coverage and resulted in loss of fragment-end coordinates critical for fragmentomic analyses

**Figure 2:**
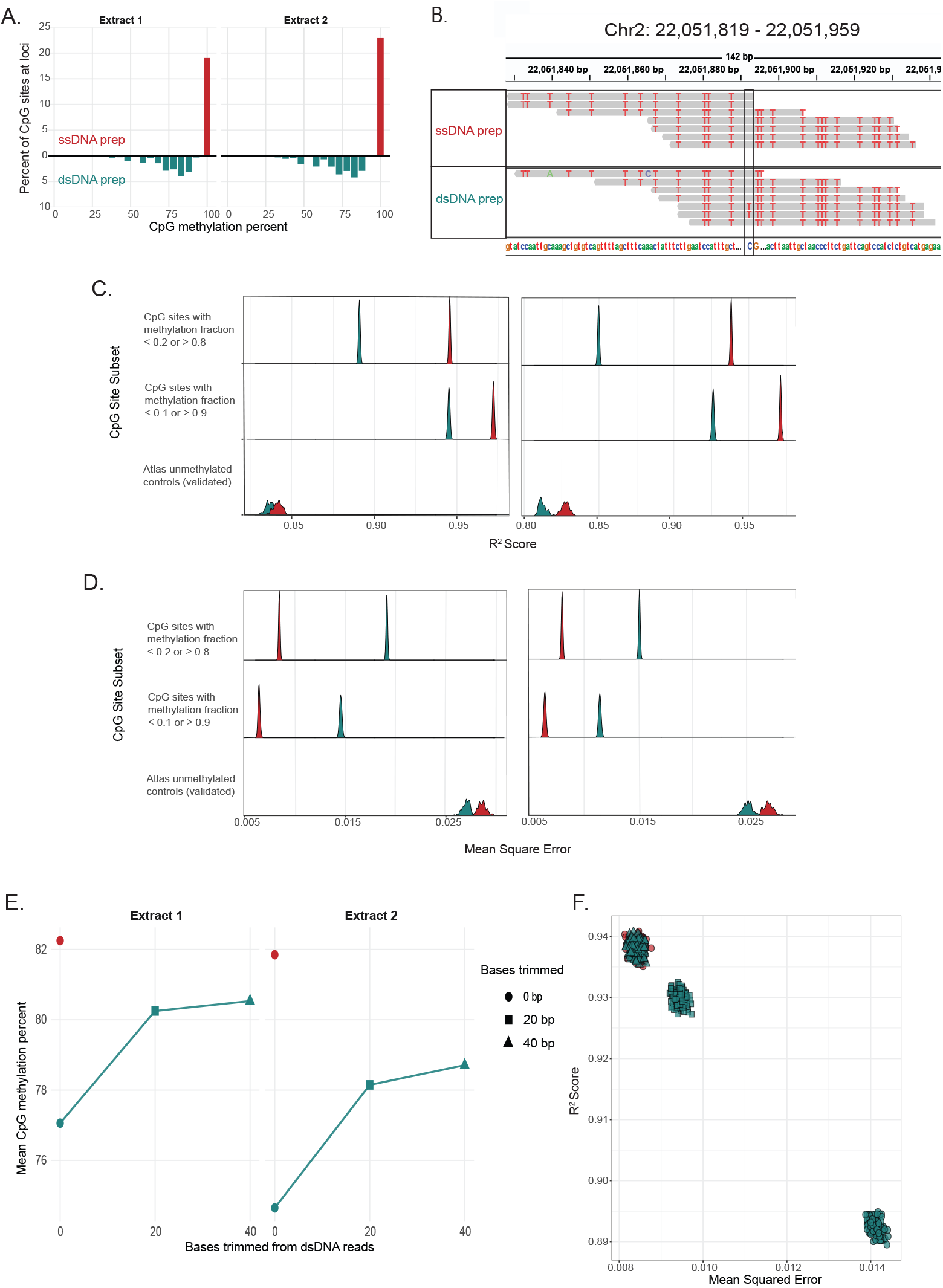
Consequences of library preparation for CpG methylation distributions and tissue-of-origin (TOO) inference. A. Difference density plots of CpG methylation proportions (ssDNA minus dsDNA) for two cfDNA extracts. Values above 0 indicate higher density in ssDNA libraries at that methylation level, values below 0 indicate higher density in dsDNA libraries. B. Representative IGV snapshot at a CpG site. Individual reads are shown as arrows aligned to the reference genome, with methylated CpGs marked. ssDNA libraries retain methylation calls at fragment ends whereas dsDNA libraries show terminal losses. C. NNLS concordance (R^2^) between cfDNA libraries and a plasma-relevant reference atlas of granulocytes, monocytes, CD4 and CD8 T cells, liver hepatocytes, kidney epithelium, and lung endothelium. Each ridge represents the density of 100 iterations of a 70% train and 30% test result. D. NNLS error (mean squared error, MSE) for the same analyses shown in C. E. Average CpG methylation (%) across dsDNA and ssDNA libraries after progressive 3’-end trimming of dsDNA reads (0, 20, and 40 bp). F. Relationship between NNLS concordance (R^2^) and error (MSE) for the informative CpG subset (<0.2 or >0.8 fraction methylation) comparing ssDNA 0 bp trimmed library to dsDNA library trimmed 0, 20, or 40 bp. Each point corresponds to one of 100 iterations from the analysis described in C–D.

### Discussion

Subtle differences in NGS library preparation chemistry directly alter the methylation landscape of cfDNA. Both ssDNA and dsDNA libraries capture the essential features of the cfDNA methylome, including the canonical dip at nucleosome dyads and phased methylation across nucleosomes that arise from restricted DNA methyltransferase access within the nucleosome core. However, dsDNA libraries show a uniform decrease in CpG methylation and a strand-oriented downstream loss resulting from the incorporation of unmethylated cytosines during end-repair fill-in. This process not only replaces the native 3’ terminal sequence of cfDNA fragments but also modifies the methylation state at those positions, altering the true biochemical composition of the library. By comparing ssDNA and dsDNA preparations generated from the same cfDNA extracts, we distinguish end-repair–dependent chemical alterations from methylation patterns that more faithfully reflect the native state of cfDNA fragments. This distinction is critical because cfDNA methylomes are now widely used to infer tissue of origin and to study chromatin organization in health and disease.

Bioinformatic end trimming provides a practical means to reduce end-repair–induced methylation changes, but it comes with inherent tradeoffs in data yield and interpretability. Consistent with prior methyl-seq studies on genomic DNA, trimming several bases from the 3’ end of the template read decreases the influence of end-repair synthesis on methylation profiles by removing bias-prone termini [21]. In cfDNA, where fragmentation produces longer overhangs and a modal fragment size of ∼167 bp, more extensive trimming is required to adjust dsDNA methylation and tissue-of-origin results toward those observed with ssDNA libraries. The magnitude of this adjustment varies across samples: one cfDNA extract in this study showed a lower dsDNA methylation baseline even before trimming, potentially reflecting biological differences or sample-specific degradation during collection or storage. This variability suggests that fixed trimming thresholds may not fully normalize across studies or sample types, as illustrated by the comparison between Fig. 2F and Supplementary Fig. 1. Alternative approaches are also emerging. A recent report described a duplex-based computational correction method (JEEPERS) that reconstructs cfDNA molecules from complementary strands to detect and correct end-repair–associated methylation changes in silico[22]. While such methods can recover 3’-end methylation without trimming, they rely on duplex sequencing, which inherently reduces library complexity because complete duplex recovery is rarely achieved. Collectively, these findings demonstrate that end repair consistently alters the biochemical composition of dsDNA libraries and that both trimming and duplex-aware correction can mitigate these effects to varying degrees, each with associated tradeoffs in sequencing efficiency and data completeness.

Together, these findings refine our understanding of how technical and biological factors jointly shape cfDNA methylation landscapes. By directly comparing ssDNA and dsDNA preparations from identical cfDNA extracts, we reveal the extent to which end-repair modifies the biochemical composition of methyl-seq libraries. This comparison clarifies why differences in library chemistry produce measurable shifts in methylation and fragmentomic signals, particularly in cfDNA and other degraded DNA sources prone to long overhangs, such as FFPE and ancient DNA. As cfDNA methylation profiling becomes increasingly central to early cancer detection and minimal residual disease monitoring, the choice of library preparation method will have a direct impact on data accuracy and interpretability. More broadly, integrating fragmentomic and methylomic analyses at this resolution will improve models of cfDNA generation, degradation, and turnover, informing both assay design and biological interpretation. Ultimately, while both dsDNA and ssDNA libraries recover the core fragmentomic features of cfDNA, ssDNA preparation eliminates end-repair–induced methylation changes and preserves native fragment termini, providing a more accurate framework for cfDNA methylation profiling.

## Abbreviations

cfDNA: cell-free DNA
TOO: tissue of origin
ssDNA: single-stranded DNA
dsDNA: double-stranded DNA
CpG: cytosine–phosphate–guanine dinucleotide
DNASE1L3: deoxyribonuclease 1 like 3
5mC: 5-methylcytosine
hg38: human genome build 38
GEO: Gene Expression Omnibus
NNLS: non-negative least squares
MSE: mean squared error
IGV: Integrative Genomics Viewer
R^2^: coefficient of determination
FFPE: formalin-fixed, paraffin-embedded
R1: read 1
R2: read 2
SRA: Sequence Read Archive
HHS: U.S. Department of Health and Human Services
CFR: Code of Federal Regulations
NGS: next-generation sequencing
Q20: Phred quality score of 20

## Declarations

### Ethics approval and consent to participate

This work is not considered human subjects research under the HHS human subjects regulations (45 CFR Part 46). Whole blood from healthy donors was commercially purchased from Stanford Blood Center, Palo Alto, CA. Donors were deidentified, no biographic or clinical information was provided to Claret Bioscience LLC.

### Consent for Publication

Not applicable

### Data and Materials Availability

The raw fastqs for the libraries used in this analysis are available for download. They can be found under the SRA submission: PRJNA1338729. The code generated from this study is available at https://github.com/tgroth97/cfdna-methylseq-comparison.

### Authors’ Contributions

VR and CJT conceptualized the study. TG, AM, and CJT designed the study. AM and RT performed the experimentation and acquired the data. TG, AM, VR, RT, and CJT analyzed the data and interpreted the results. TG and CJT performed figure generation. TG and CJT drafted and revised the manuscript with input from AM, VR, RT. TG, AM, VR, RT, CJT approved the final manuscript.

## Acknowledgements

Not Applicable

---

**Supplementary Table 1.**
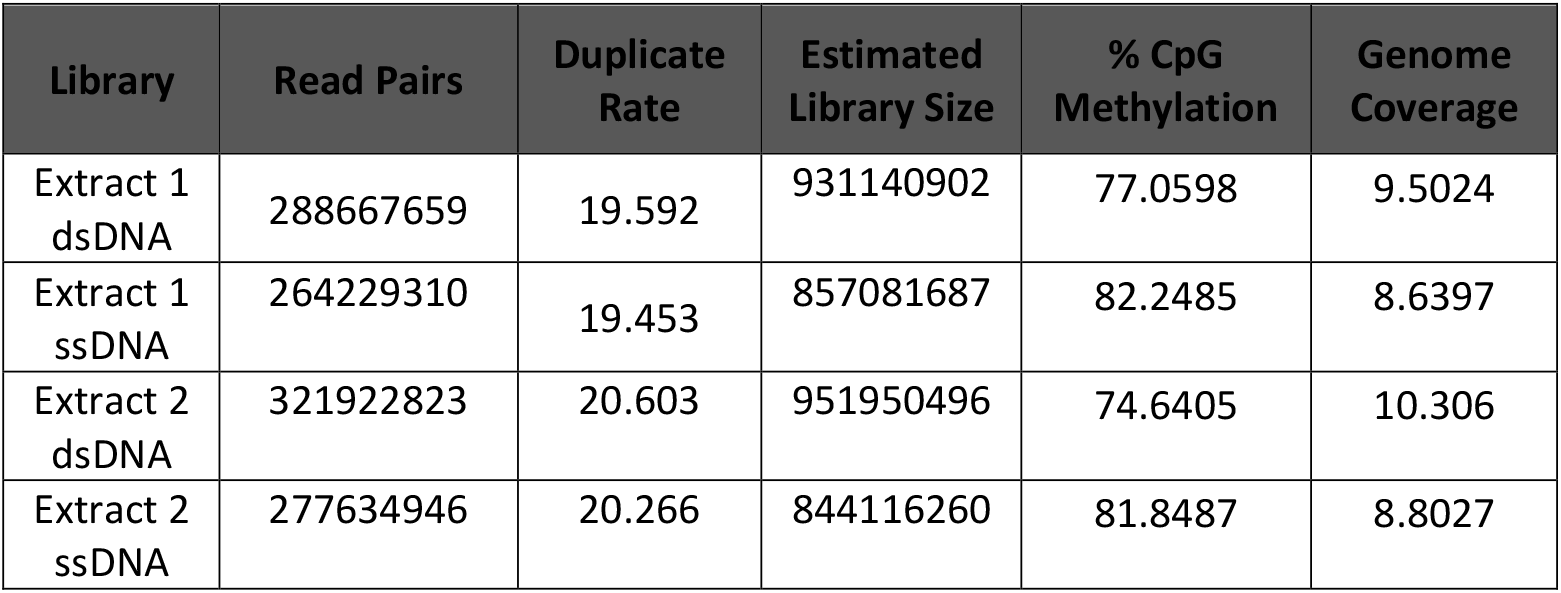
This table contains different stats regarding the libraries used in our analysis. The read pairs are calculated from the number of read pairs contained in the raw fastq files. Duplicate rate and estimated library size are output by Picard’s MarkDuplicates tool. Percent CpG Methylation was calculated from the bed file output by MethylDackel extract. Genome coverage was calculated using the samtools depth tool.

**Supplementary Table 2.**
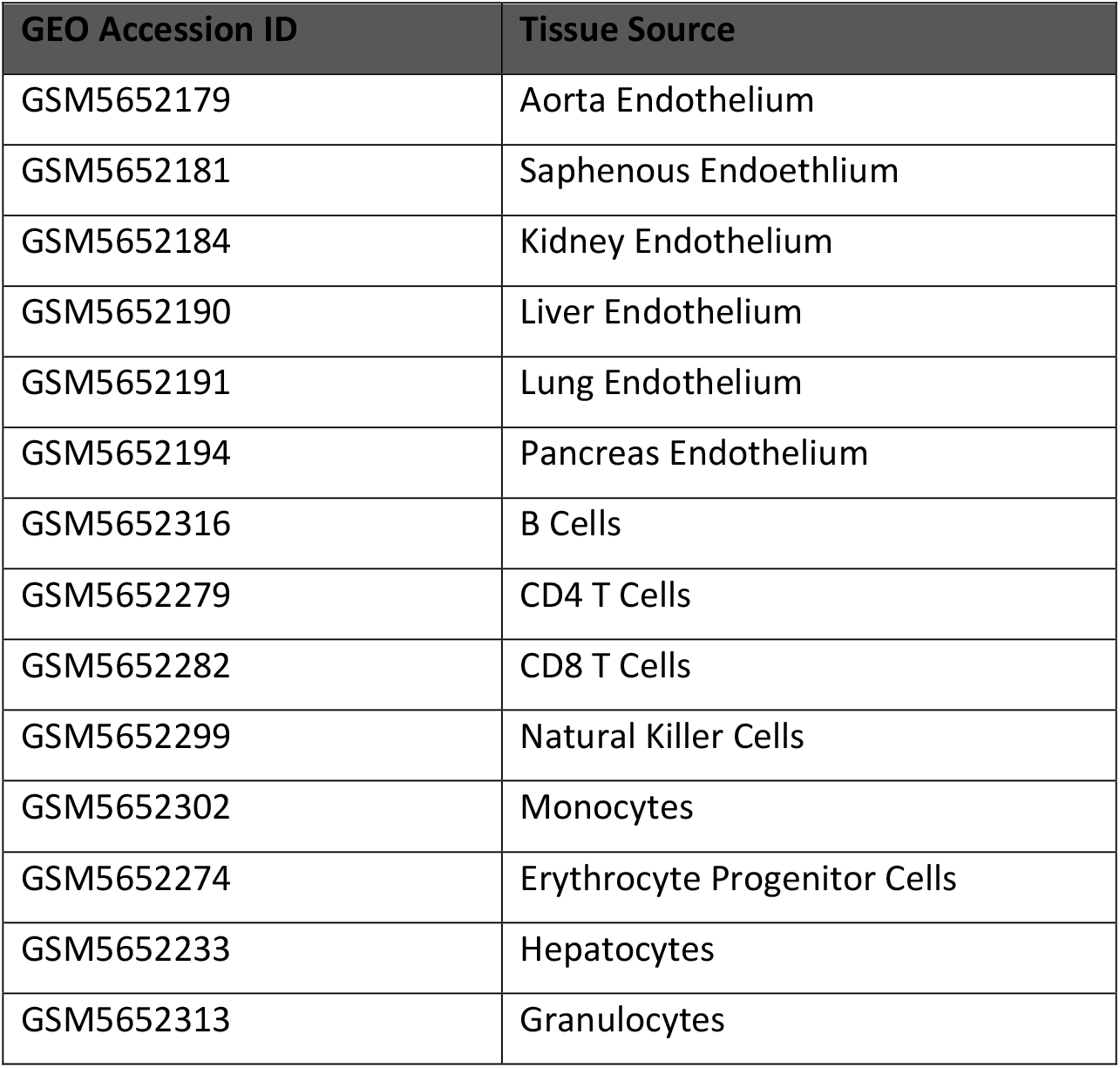
The tissues used in our literature informed cfDNA associated tissue set are seen in the table above. In Loyfer et al, they identified which tissue types contributed to cfDNA, and we selected tissues from their analysis that fell into those identified tissue types. The tissues they identified are as follows: vascular endothelium, lymphocytes, monocytes, erythrocyte progenitors, megakaryocytes, hepatocytes, and granulocytes. The GEO accession ID for the full project is GSE186458, and we have provided the specific GEO IDs for each tissue we retrieved. For each tissue sample, we downloaded the hg38 BigWig file.

**Supplementary Figure 1.**
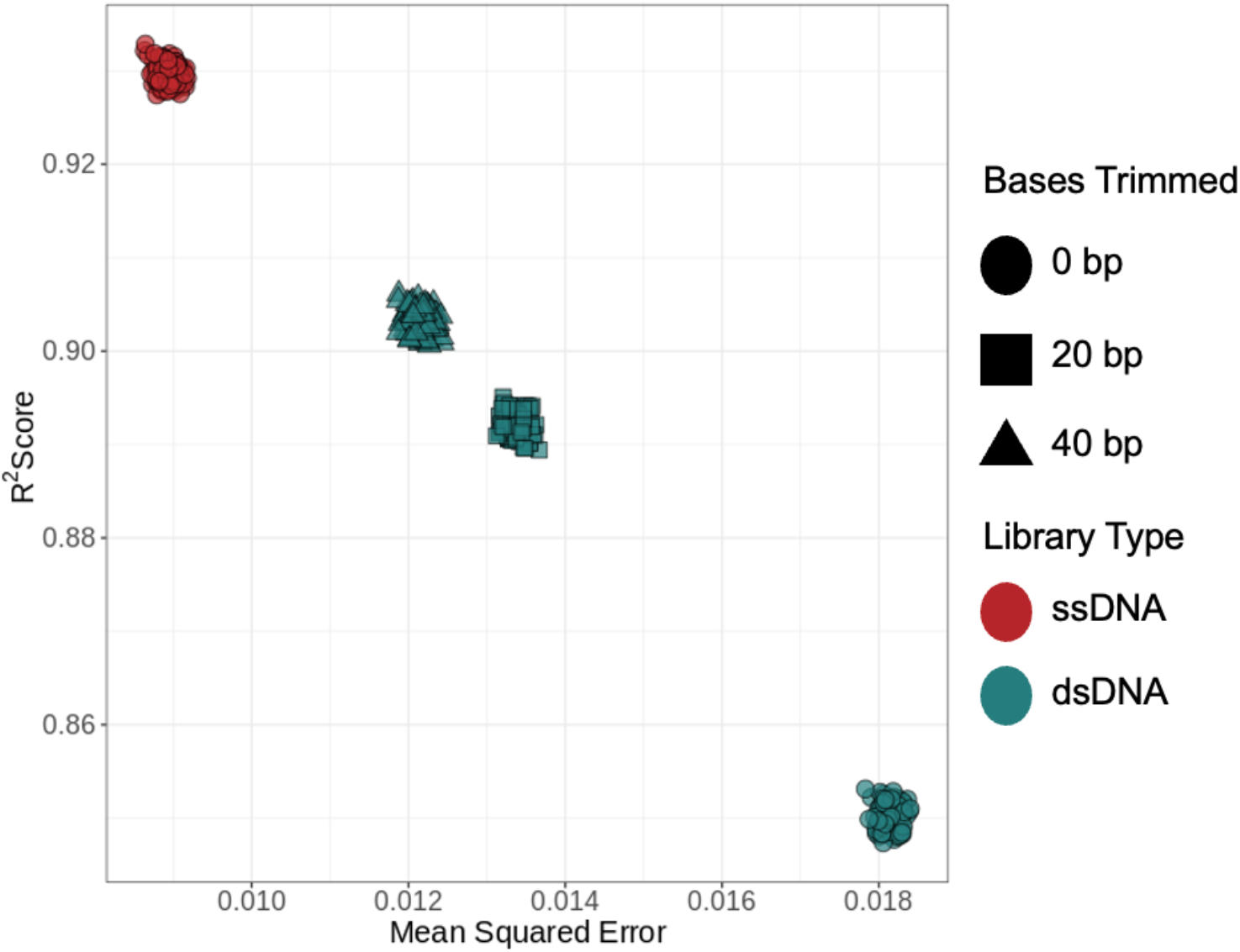
Relationship between NNLS concordance (R^2^) and error (MSE) for the informative CpG subset (<0.2 or >0.8 fraction methylation) comparing ssDNA 0 bp trimmed library to dsDNA library trimmed 0, 20, or 40 bp, using the other matched library pair.

