## Supplemental for "End-repair causes methylation underestimation in cell-free DNA sequencing libraries"

| **Library** | **Read Pairs** | **Duplicate Rate** | **Estimated Library Size** | **% CpG Methylation** | **Genome Coverage** |
| --- | --- | --- | --- | --- | --- |
| Extract 1 dsDNA | 288667659 | 19.592 | 931140902 | 77.0598 | 9.5024 |
| Extract 1 ssDNA | 264229310 | 19.453 | 857081687 | 82.2485 | 8.6397 |
| Extract 2 dsDNA | 321922823 | 20.603 | 951950496 | 74.6405 | 10.306 |
| Extract 2 ssDNA | 277634946 | 20.266 | 844116260 | 81.8487 | 8.8027 |

**Supplementary Table 1**

This table contains different stats regarding the libraries used in our analysis. The read pairs are calculated from the number of read pairs contained in the raw fastq files. Duplicate rate and estimated library size are output by Picard’s MarkDuplicates tool. Percent CpG Methylation was calculated from the bed file output by MethylDackel extract. Genome coverage was calculated using the samtools depth tool.

| **GEO Accession ID** | **Tissue Source** |
| --- | --- |
| GSM5652179 | Aorta Endothelium |
| GSM5652181 | Saphenous Endoethlium |
| GSM5652184 | Kidney Endothelium |
| GSM5652190 | Liver Endothelium |
| GSM5652191 | Lung Endothelium |
| GSM5652194 | Pancreas Endothelium |
| GSM5652316 | B Cells |
| GSM5652279 | CD4 T Cells |
| GSM5652282 | CD8 T Cells |
| GSM5652299 | Natural Killer Cells |
| GSM5652302 | Monocytes |
| GSM5652274 | Erythrocyte Progenitor Cells |
| GSM5652233 | Hepatocytes |
| GSM5652313 | Granulocytes |

**Supplementary Table 2**

The tissues used in our literature informed cfDNA associated tissue set are seen in the table above. In Loyfer et al, they identified which tissue types contributed to cfDNA, and we selected tissues from their analysis that fell into those identified tissue types. The tissues they identified are as follows: vascular endothelium, lymphocytes, monocytes, erythrocyte progenitors, megakaryocytes, hepatocytes, and granulocytes. The GEO accession ID for the full project is GSE186458, and we have provided the specific GEO IDs for each tissue we retrieved. For each tissue sample, we downloaded the hg38 BigWig file.

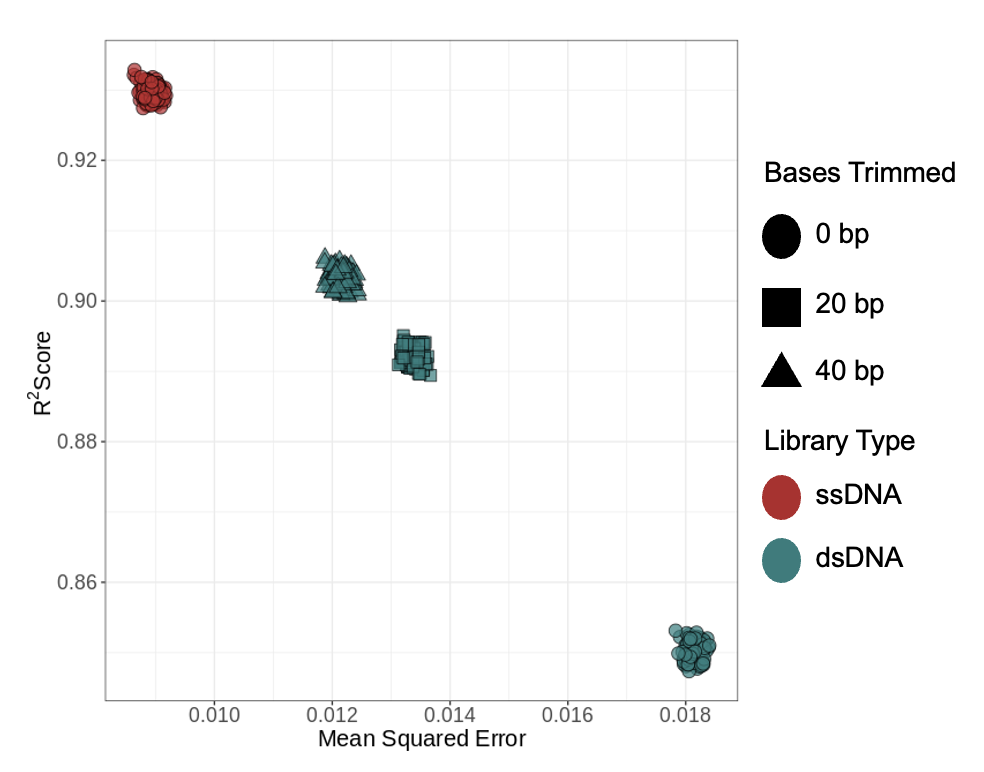

**Supplementary Figure 1**

Relationship between NNLS concordance (R²) and error (MSE) for the informative CpG subset (<0.2 or >0.8 fraction methylation) comparing ssDNA 0 bp trimmed library to dsDNA library trimmed 0, 20, or 40 bp, using the other matched library pair.
